# Dynamic Effects of Ventral Hippocampal NRG3/ERBB4 Signaling on Nicotine Withdrawal-Induced Responses

**DOI:** 10.1101/2023.01.17.524432

**Authors:** Miranda L. Fisher, Emily R. Prantzalos, Bernadette O’Donovan, Tanner Anderson, Pabitra K. Sahoo, Jeffery L. Twiss, Pavel I. Ortinski, Jill R. Turner

## Abstract

Tobacco smoking remains a leading cause of preventable death in the United States, with a less than 5% success rate for smokers attempting to quit. High relapse rates have been linked to several genetic factors, indicating that the mechanistic relationship between genes and drugs of abuse is a valuable avenue for the development of novel smoking cessation therapies. For example, various single nucleotide polymorphisms (SNPs) in the gene for neuregulin 3 (*NRG3*) and its cognate receptor, the receptor tyrosine-protein kinase erbB-4 (*ERBB4)*, have been linked to nicotine addiction. Our lab has previously shown that ERBB4 plays a role in anxiety-like behavior during nicotine withdrawal (WD); however, the neuronal mechanisms and circuit-specific effects of NRG3-ERBB4 signaling during nicotine and WD are unknown. The present study utilizes genetic, biochemical, and functional approaches to examine the anxiety-related behavioral and functional role of NRG3-ERBB4 signaling, specifically in the ventral hippocampus (VH). We report that 24hWD from nicotine is associated with altered synaptic expression of VH NRG3 and ERBB4, and genetic disruption of VH *ErbB4* leads to an elimination of anxiety-like behaviors induced during 24hWD. Moreover, we observed attenuation of GABAergic transmission as well as alterations in Ca^2+^-dependent network activity in the ventral CA1 area of VH *ErbB4* knock-down mice during 24hWD. Our findings further highlight contributions of the NRG3-ERBB4 signaling pathway to anxiety-related behaviors seen during nicotine WD.

## Introduction

Nicotine addiction impacts 1.2 billion people worldwide, with more people addicted to nicotine than any other drug [1]. Abstinence from chronic nicotine use results in both cognitive and affective withdrawal (WD) symptoms, which can be observed just a few hours after discontinuation of nicotine use [2] and are suggested to be the predominant factors in driving relapse to cigarette smoking [3]. Supporting data link hippocampal function with nicotine WD-induced phenotypes in both humans [4-8] and rodents [9-12]. However, mounting evidence suggests that the hippocampus is not a homogenous structure, but instead, it can be divided into dorsal and ventral regions, each mediating different behaviors [13]. Our lab has previously reported that these subregional functional differences correspond with distinct WD phenotypes. We found that cAMP response element-binding protein (CREB) activity, specifically in the ventral hippocampus (VH), mediates anxiety-like behaviors in mice undergoing 24hWD, whereas dorsal hippocampal (DH) CREB activity mediates cognitive effects [14]. Furthermore, to elucidate potential CREB target genes underlying these phenotypes, we evaluated CREB binding genome-wide following chronic nicotine exposure and WD using chromatin immunoprecipitation and whole-genome sequencing. These experiments showed that CREB is highly enriched at the promoter for the Neuregulin-3 (*Nrg3*) gene following chronic nicotine and WD in the hippocampus [15].

NRG3 is a neuronal-enriched member of the epidermal growth factor-like (EGF-like) family of Neuregulins 1-6. NRG3’s expression is limited to the CNS, where its EGF-like domain binds exclusively to receptor tyrosine-protein kinase erbB-4 (ERBB4) receptors [16] enriched in neuronal post-synaptic densities (PSD) of inhibitory interneurons [17-19]. In situ hybridization studies show that *Nrg3* and *ErbB4* have the highest expression in cortical and hippocampal regions [16], where their interactions play pleiotropic roles in brain development and plasticity. NRG3 was identified as a chemoattractive factor regulating GABAergic interneuron migration through its interaction with ERBB4 in the developing brain [20]. NRG3-ERBB4’s involvement in the assembly and maturation of inhibitory circuitry is particularly noteworthy due to its association with a wide variety of neurodevelopmental and neuropsychiatric disorders [21]. Less is known about this pathway’s function in the adult brain, but it is speculated to remain involved in activity-dependent synaptic formation and maintenance. Addictive drugs are known to cause persistent restructuring of several different neuronal subtypes resulting in long-term changes in synaptic plasticity. We have previously demonstrated that ERBB4 activation is necessary for nicotine-induced plasticity in the orbitofrontal cortex, a region associated with impulse control [22]. Additionally, our previous investigations revealed that ERBB4 antagonism attenuates anxiety-like behavior during chronic nicotine WD [15]. However, circuit-specific effects of NRG3-ERBB4 signaling on the affective measures of prolonged nicotine exposure and WD have yet to be previously determined.

Genetic association studies from our lab and collaborators indicate a significant association of multiple *NRG3* and *ERBB4* single nucleotide polymorphisms (SNPs) with smoking cessation outcomes [15,23]. While there is persuasive evidence for the role of NRG3-ERBB4 signaling in nicotine dependence, the precise activity of these signaling molecules and the neural adaptations they regulate during WD from nicotine is unknown. Therefore, this study aims to systematically investigate the VH ERBB4 signaling during chronic nicotine and WD. We found that selective deletion of VH *ErbB4* attenuates anxiety-like behavior during nicotine WD. This anxiolytic effect was accompanied by reductions in inhibitory synaptic transmission and alterations in network activity of ventral CA1 pyramidal neurons.

## Results

### 24hWD from nicotine induces Nrg3 transcription in the ventral hippocampus

NRG3 and ERBB4 are highly enriched at excitatory synapses onto interneurons within the VH [33]. To determine nicotine effects on NRG3 and ErbB4 expression in this structure, wild-type *ErbB4*-floxed mice were treated with intermittent saline or nicotine (12 mg/kg/day) and were subjected to 24h or 1week of WD. Ventral hippocampal tissues collected from treated mice were used for mRNA and protein analysis.

RTqPCR analysis of wild type *ErbB4*-floxed mice revealed that VH *Nrg3* mRNA levels are increased at the 24hWD time point and return to baseline by 1week, compared to chronic nicotine treated mice, suggesting alterations in *Nrg3* synthesis occur early during nicotine abstinence and return to baseline within 1week of WD (F(3,33)=3.653, p=0.0223; one-way ANOVA; NIC versus 24hWD: p=0.0117; NIC versus 1week WD: p=0.3933; post hoc analyses) (Fig.1Ai). No differences were observed in *ErbB4* mRNA levels during saline, nicotine, or either WD time point (F(3,33)=0.8802, p=0.4614; one-way ANOVA) (Fig.1Aii). Western blot analyses of the 75kDa band detected using the anti-NRG3 antibody revealed an increase in synaptosomal NRG3 protein during 24hWD, compared to saline control mice (F(2,22)=4.923, p=0.0171; one-way ANOVA; SAL versus 24hWD: p=0.0128; post hoc analyses) (Fig.1Bi). Synaptosomal fractions immunoblotted with the affinity-purified anti-ERBB4 antibody yielded major bands at 180, 120, and 80kDa [34]. The full-length 180kDa band (F(2,22)=6.974, p=0.0045; one-way ANOVA; NIC vs. 24hWD: p=0.0036) and the 120kDa band (F(2,21)=7.566, p=0.0034; one-way ANOVA; SAL versus 24hWD: p=0.0024) of ERBB4 were significantly increased during 24hWD, compared to their chronic nicotine or saline-treated counterparts (Fig.1Bii,iii). A qualitatively similar effect observed at the 80kDa band was not significant (Fig.1Biv) (F(2,20)=1.855, p>0.05).

**Figure 1.**
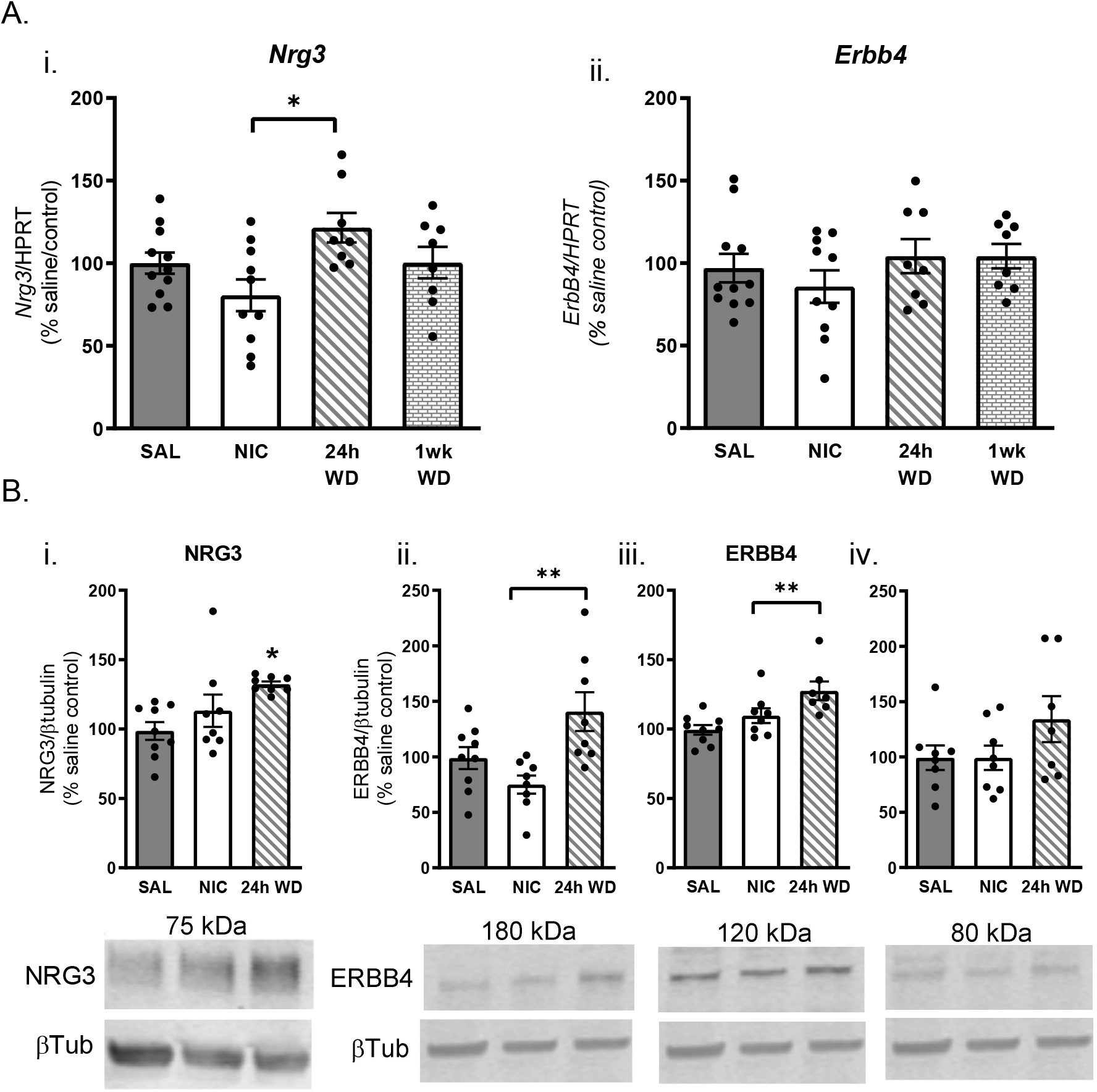
24hWD increases VH expression of NRG3 and ERBB4. (A) mRNA quantification of (i) *Nrg3* and (ii) *ErbB4*. (B) densitometry analysis of (i) synaptosomal NRG3 (75kDa band) protein, and synaptosomal ERBB4 protein at (ii) 180kDa, (iii) 120kDa, and (iv) 80kDa bands in treated ventral hippocampal murine tissue. The same Beta-tubulin control for ERBB4 analysis is shown in Bii, iii, and iv, as the analysis was conducted on a single gel. N=8-10 per group (*p<0.05, **p<0.01).

### Decreased ErbB4 within the VH attenuates 24hWD-induced transcription of Nrg3 mRNA

To target ErbB4 signaling in the VH, we performed stereotaxic microinjections of AAV-CRE, or AAV-RFP control virus, into the VH of the *ErbB4*-floxed mouse line. RFP viral expression in the VH is shown in the *Supplementary Information* to demonstrate appropriate targeting of the viral construct (Fig.S1A). These animals were treated chronically with saline or nicotine and underwent 24h of nicotine withdrawal before behavioral testing (Fig.2A). RTqPCR analysis of VH tissue demonstrated CRE recombinase expression was significantly higher in CRE-injected mice, compared to RFP-controls (t(70)=6.059 p<0.0001; unpaired t-test) (Fig.S1B). This increase in CRE recombinase expression resulted in a significant interaction and main effect of genotype and reduced *ErbB4* mRNA levels to ∼60% of that of RFP SAL mice (main effect of genotype F(1,66)=121.0, p<0.0001; two-way ANOVA; RFP SAL versus CRE SAL: p<0.0001, CRE NIC: p<0.0001, CRE 24hWD: p<0.0001, Post-hoc analyses) (Fig.2B). Furthermore, *Nrg3* mRNA expression analyses showed an interaction and main effect of genotype, with increased *Nrg3* expression in RFP 24hWD mice compared to their NIC treated counterparts (main effect of genotype F(1,66)=12.37, p<0.005; interaction F(2,66)=10.90, p<0.0001; two-way ANOVA; RFP SAL versus RFP 24hWD: p=0.0018, post hoc analyses) (Fig.2C). This increase in *Nrg3* mRNA was not observed in CRE-injected mice (no treatment effect, F(2,31=1.982, p>0.05; one-way ANOVA) (Fig.2C).

**Figure 2.**
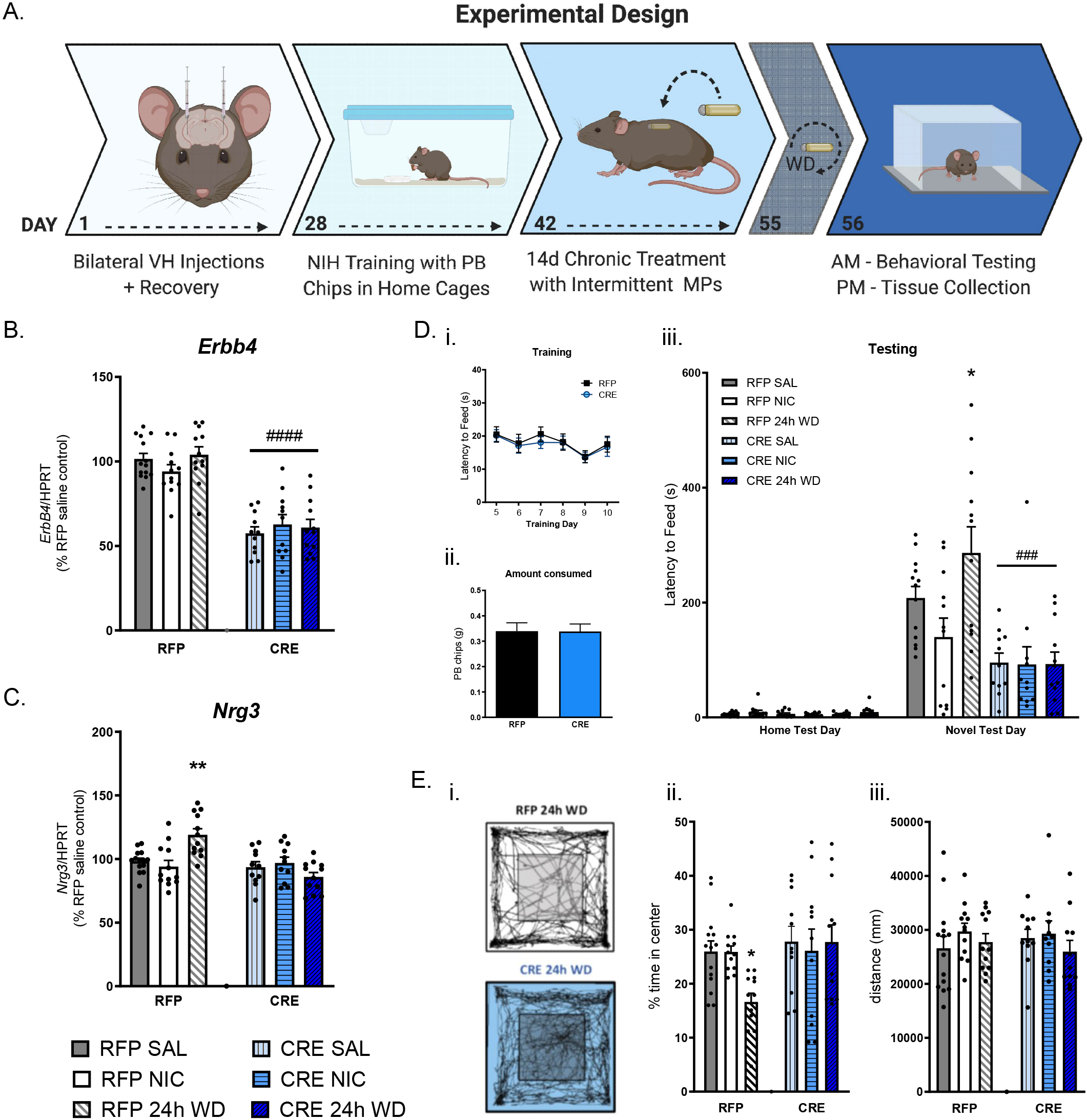
Gene expression analysis of VH *ErbB4* knock-down and consequent attenuation of nicotine WD-induced anxiety-like behavior. (A) Experimental Design. (B) Quantification of *ErbB4* mRNA in CRE-injected mice compared to RFP-injected controls. (C) Quantification of *Nrg3* mRNA in CRE-injected mice compared to RFP-injected controls. n=11-14 per group. (*p<0.05, treatment effect; ##p<0.01, ####p<0.0001, viral effect). (D) Novelty-induced hypophagia (NIH) test. (i) Latency to feed in RFP and CRE-injected animals during NIH training prior to treatment. (ii) Amount of peanut butter chips (PB) consumed in RFP and CRE-injected animals on NIH training day 10. (iii) Latency to feed in treated, RFP and CRE-injected animals in home cage (Home Test Day) and novel environment (Novel Test Day). (E) Open Field exploratory test. (i) Representative activity traces in RFP and CRE-injected mice undergoing 24hWD. (ii) Measurement of percent time spent in the center arena of treated RFP- and CRE-injected mice. (iii) Locomotor activity of RFP- and CRE-injected mice in arena. N=11-14 per group. (*p<0.05, treatment effect; ###p<0.001, viral effect).

### VH ErbB4 knock-down blocks anxiogenic behavior measured in the NIH test

We next performed behavioral analyses to evaluate the influences of VH *ErbB4* KD on anxiety-like behaviors. The NIH test is a well-validated measure for VH-dependent anxiety-related behaviors that is sensitive to acute anxiolytics and chronic antidepressants [35]. No genotype effects were observed during training in latency to consume prior to treating mice with saline or nicotine (t(5)=2.162, p>0.05; paired t-test) (Fig.2Di) or amount consumed (t(70)=0.02770, p>0.05; unpaired t-test) (Fig.2Dii). Mice then underwent two weeks of chronic treatment of saline or nicotine via osmotic minipumps. To confirm there were no appetitive effects of chronic nicotine treatment, mice were presented with peanut butter chips (PB) in their home cage, and latency was measured (Home Test Day). No significant difference between groups was observed (F(5,66)=1.072, p=0.3940; one way ANOVA) (Fig.2Diii). After Home Test Day, osmotic minipump removal and sham surgeries were performed to induce WD. After 24hWD, mice were placed in a novel environment (Novel Test Day) and again latency to feed was measured, displaying an interaction and main effect of treatment, and day (main effect of day, F(1, 131)=147.6, p<0.0001; main effect of treatment F(5, 131)=7.281, p<0.0001; interaction F(5, 131)=7.458, p<0.0001) (Fig.2Diii). RFP mice undergoing 24hWD displayed a significant increase in latency to feed on novel test day compared to controls (RFP SAL versus RFP 24hWD: p=0.0302; post hoc analyses), while CRE-injected animals showed a significant reduction in latency to feed across all treatment groups, compared to RFP SAL control mice (RFP SAL versus CRE SAL: p=0.0009; CRE NIC: p=0.0006; CRE 24hWD: p=0.0005; post-hoc analyses) (Fig.2Diii). No sex differences were observed.

After NIH testing was conducted, mice were placed in their home cages for a 1h recovery period before the Open Field exploration test was run that afternoon. Representative traces show activity in the Open Field arena of RFP- and CRE-injected 24hWD groups (Fig.2Ei). Results from this test indicated a significant treatment effect (F(1,66)=5.148, p<0.05; two-way ANOVA) (Fig.2Eii). Control mice undergoing 24hWD (RFP 24hWD) displayed a significant decrease in time spent in the center arena, compared to SAL controls (p=0.0290), indicative of an anxiogenic response to a novel environment during WD (Fig.2Eii). Interestingly, in CRE-injected VH *ErbB4* KD mice, this treatment effect was undetectable, with no significant differences between SAL and 24hWD groups (RFP SAL versus RFP 24hWD: p=.9994; post hoc analyses) (Fig.2Eii). No locomotor deficits were observed between genotypes or treatments (no interaction F(2,64)=0.4727, p>0.05; two-way ANOVA) (Fig.2Eiii). Collectively, findings from both behavioral tests suggest that disruption of NRG3-ERBB4 activity in the VH attenuates prominent anxiogenic effects induced during 24hWD.

### ErbB4 mRNA expression predominates in the CA1 area of the ventral hippocampus

The hippocampus has a very well-defined architecture consisting of populations of excitatory principal neurons assembled into distinct regions: the dentate gyrus (DG), and areas CA3-CA1. These areas form what is known as the trisynaptic circuit, an information flow beginning with cortical inputs from the entorhinal cortex (EC), which carry higher-order spatial and contextual information, synapsing onto DG granule cells via perforant path fibers. Mossy fibers from DG granule cells project to pyramidal neurons of the CA3 which form schaffer collaterals innervating CA1 pyramidal cells. Axons of CA1 pyramidal neurons in turn target both intra-hippocampal and extra-hippocampal territories [36]. It is well-accepted that the different circuit components along this pathway (DG, CA3, CA1) contribute to unique aspects of memory and emotional processing [37]. Therefore, circuit-specific expression and functional patterns of NRG3 and ERBB4 may provide insights into how this signaling pathway modulates circuit-level events underlying affective behaviors during nicotine WD. To this end, we used smFISH to visualize and quantify individual *Nrg3* and *ErbB4* mRNA puncta signals within the VH subregions. In Figure 3Ai, a representative 10x image of the VH illustrates patterns of peri-nuclear expression (Fig.3Ai). Images of the subregions DG, CA3, and CA1 of the VH, highlighted in yellow in the right (Fig.3Aii), were taken with a 63x oil objective to observe expression patterns of *Nrg3* (Fig.3B) and *ErbB4* mRNAs (Fig.3D). Quantitative analysis using ImageJ showed consistent expression of *Nrg3* mRNA within the DG, CA3, and CA1 areas of the VH, with no significant differences between regions (F(2,6)=1.659, p>0.05; one way ANOVA) (Fig.3C). Conversely, quantification of *ErbB4* mRNA showed a significant difference in signal between the CA1 and DG (F(2,6)=8.977, P<0.05; one way ANOVA), with the highest expression of *ErbB4* mRNA present in the CA1 area of the VH (DG versus CA1, p=0.0138) (Fig.3E).

**Figure 3.**
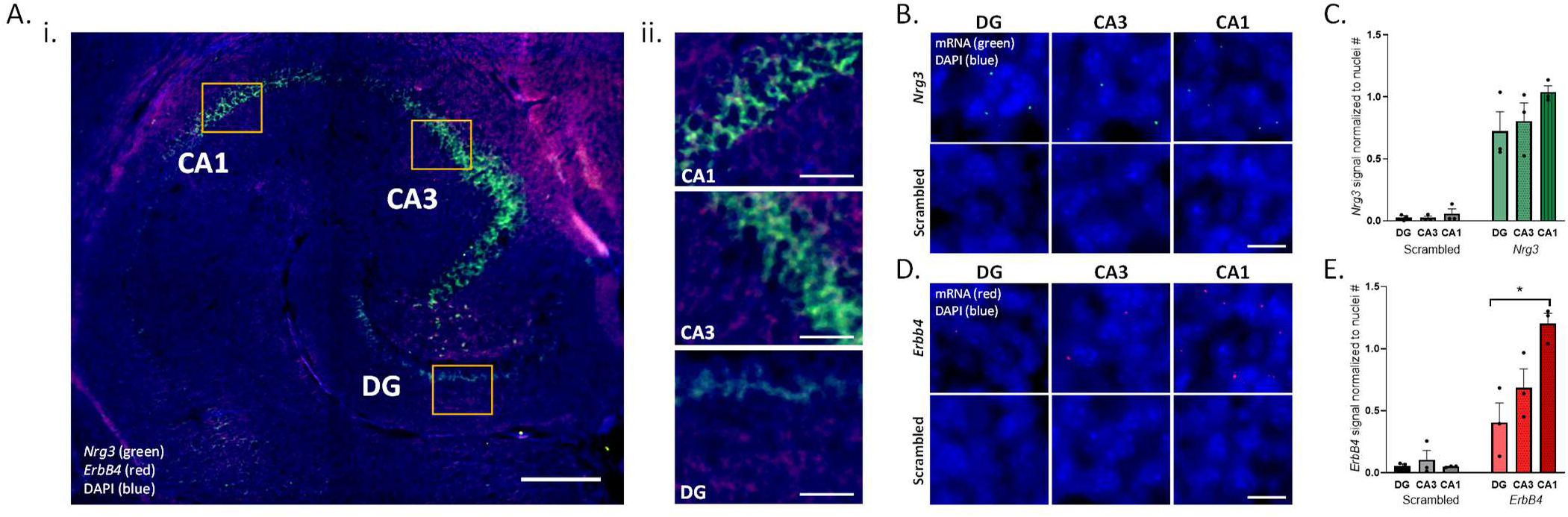
*Nrg3* and *ErbB4* mRNA expression predominates in the CA1 region of the VH. Representative horizontal 30 μm VH sections from untreated mice, incubated with either Stellaris probe against *Nrg3* or *ErbB4* mRNAs or scrambled probes. Background subtractions are based on scrambled control images. Representative 10x image of *Nrg3* (green) and *ErbB4* (red) mRNA labeled with Stellaris RNA FISH probes, and DAPI stain (blue) in (i) the VH (Scale bar 300 μm), and (ii) subareas DG, CA3, and CA1 (Scale bar 50 μm). Representative 63x oil immersion images *Nrg3* mRNA puncta (green) and DAPI (blue) in the DG, CA3, CA1 of VH (Scale bar 20 μm). (C) Quantification of *Nrg3* mRNA normalized to nuclei number. (D) Representative 63x oil immersion images of *ErbB4* mRNA puncta (green) and DAPI (blue) in the DG, CA3, and CA1 of VH. (E) Quantification of *ErbB4* mRNA normalized to nuclei number (scale bar 20 μm). N=3 per group. (*p<0.05).

### Ventral hippocampal ErbB4 knock-down reduces spontaneous IPSC and miniature IPSC frequencies in the ventral CA1

Given its central output role and high NRG3/ERBB4 expression, we next investigated the functional impact of ERBB4 on ventral CA1 neuronal activity during 24hWD. *ErbB4*-floxed animals underwent stereotaxic microinjections of either AAV-RFP+ GCaMP6f (control) or AAV-CRE+ GCaMP6f (*ErbB4* KD) in the VH, allowing for collection of electrophysiological and Ca^2+^ imaging recordings of CA1 pyramidal neurons from the same subject during the 24hWD condition. Using whole-cell patch-clamp electrophysiology, we first evaluated the effect of VH *ErbB4* KD on spontaneous excitatory and inhibitory postsynaptic currents (sEPSCs and sIPSCs) (Figure 4Ai/Bi), as well as action-potential independent miniature excitatory and inhibitory postsynaptic currents (mEPSCs and mIPSCs) in CA1 pyramidal cells. VH *ErbB4* KD significantly reduced sIPSC frequency in CA1 pyramidal neurons (t(17)=3.228 p<0.01; unpaired t-test) (Fig.4Aii), with no significant changes in sIPSC amplitude (t(17)=0.4036 p>0.05; unpaired t-test) (Fig.4Aiii). The mIPSC data also showed reduced frequency of synaptic events (t(11)=2.612 p<0.05; unpaired t-test) (Fig.4Aiv); however, we also observed an increase in mIPSC amplitude (t(11)=3.994 p<0.01; unpaired t-test) (Fig.4Av). No differences in either sEPSC frequency (t(16)=0.6112 p>0.05; unpaired t-test) (Fig.4Bii) or amplitude (t(17)=1.195 p>0.05; unpaired t-test) (Fig.4Biii) or mEPSC frequency (t(14)=0.9388 p>0.05; unpaired t-test) (Fig.4Biv) or amplitude (t(14)=0.6612 p>0.05; unpaired t-test) (Fig.4Bv) were observed in the KD mice undergoing 24hWD compared to controls. These findings show that in mice undergoing 24hWD, *ErbB4* deletion primarily impacts inhibitory synaptic transmission onto CA1 pyramidal neurons in the VH.

**Figure 4.**
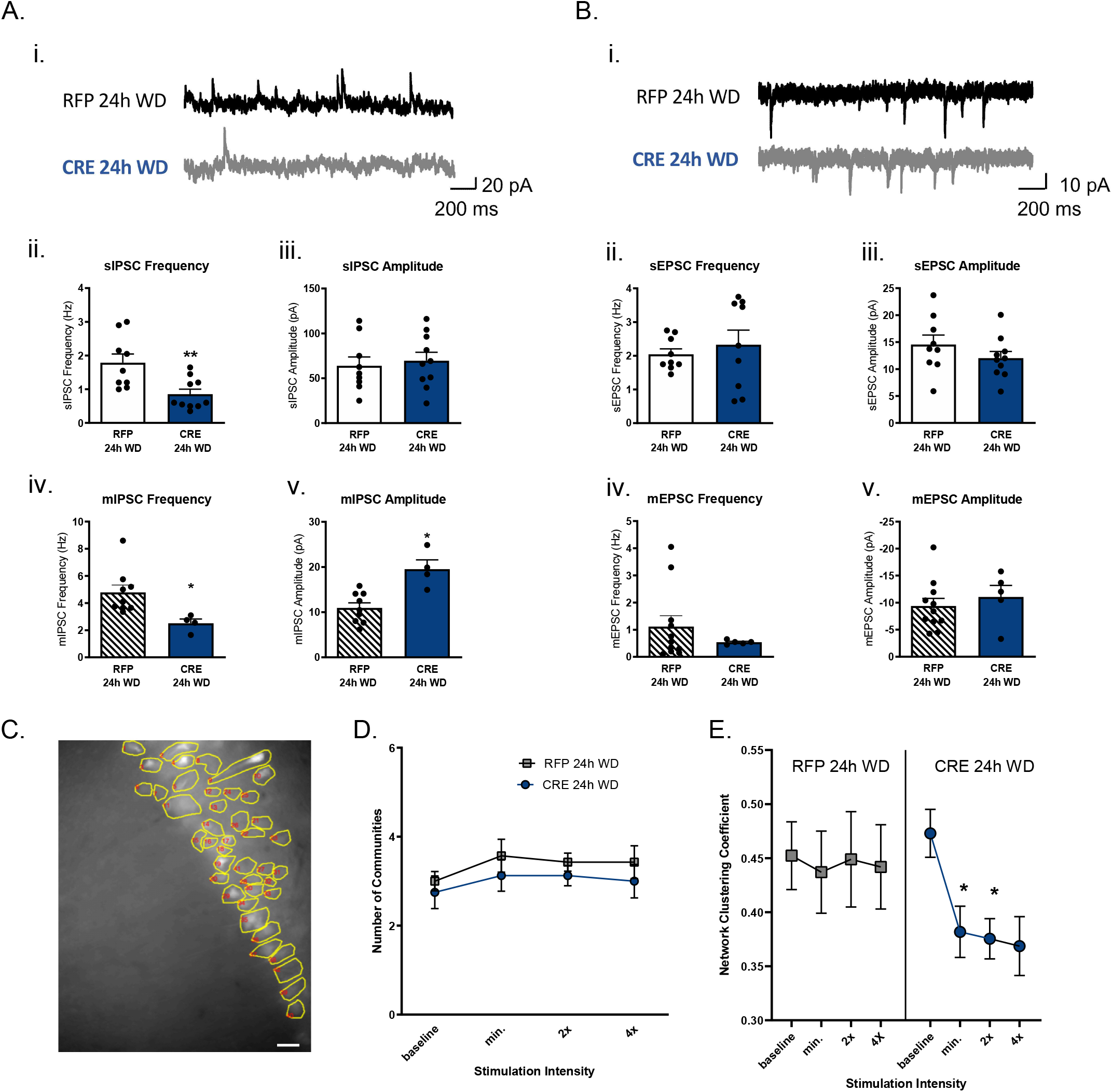
Ventral hippocampal *ErbB4* knock-down reduces spontaneous IPSC frequency and Ca^2+^- dependent network clustering during 24hWD. (A) Whole cell patch clamp recordings of spontaneous and miniature inhibitory post-synaptic currents (sIPSCs/mIPSCs). (i) Representative sIPSC traces. Quantification of sIPSC (ii) frequency and (iii) amplitude and mIPSC (iv) frequency and (v) amplitude in RFP- and CRE-injected mice undergoing 24hWD. (B) Whole cell patch clamp recordings of spontaneous and mini excitatory post-synaptic currents (sEPSCs/mEPSCs). (i) Representative sEPSC traces. Quantification of sEPSC (ii) frequency and (iii) amplitude and mEPSC (iv) frequency and (v) amplitude in RFP- and CRE-injected mice undergoing 24hWD. N=4 animals (2-3 cells per animal) (**p<0.01). (C) Representative 40x image of fluorescently GCaMP6f-labelled ventral CA1 pyramidal cells outlines as numbered ROIs (scale bar 30 microns). (D) Quantification of ventral CA1 pyramidal cell communicatees across increasing stimulation intensities in RFP- and CRE-injected mice undergoing 24hWD. (E) Quantification of network clustering coefficient across increasing stimulation intensities in RFP- and CRE-injected mice undergoing 24hWD. N=4 mice (2-3 sections per animal (*p<0.05).

### Ventral hippocampal ErbB4 knock-down impacts CA1 network activity during 24hWD

After observing alterations in inhibitory input onto individual CA1 pyramidal cells, we next queried whether these effects impacted overall network activity between genotypes using wide-field Ca^2+^ imaging. We monitored fluorescence of the hSYN-driven Ca^2+^ sensor, GCaMP6f [38], in the ventral CA1 of RFP and VH *ErbB4* KD mice undergoing 24hWD (regions of interest, ROIs, Fig.4C). Given the importance of inhibitory signaling on patterning of pyramidal cell output in the hippocampus [39], we evaluated the effect of ErbB4 KD on two measures of network segregation: community structure and network clustering. Quantification of spontaneous (baseline) and evoked Ca^2+^ transients showed no difference in the number of communities between RFP and CRE-injected mice across stimulation intensities (no interaction, F(3,39)=0.04595, p>0.05; RM two-way ANOVA) (Fig.4D). Furthermore, we found no differences in the network clustering coefficient across stimulation intensities in wild type RPF 24hWD mice (F(2.364, 14.18)=0.04814, p>0.05, RM one way ANOVA) (Fig.4E). In contrast, CRE 24hWD mice displayed a significant reduction in network clustering (F(1.996, 13.97)=5.124, p<0.05; RM one way ANOVA) at the minimum (BASELINE versus MIN. STIM: p=0.0419; post hoc analyses) and 2x (BASELINE versus 2x STIM: p=0.0497; post hoc analyses) stimulation intensities compared to baseline (Fig.4E). These data indicate that ErbB4 KD increased sensitivity of the hippocampal network to electrical stimuli at 24hWD from chronic nicotine.

## Discussion

Cumulative evidence indicates that while the positive reinforcing effects of nicotine play a crucial role in the development of nicotine dependence, negative reinforcers, such as WD symptoms, drive the maintenance of nicotine dependence similar to other stimulants [40]. For example, the ability of nicotine to alleviate the negative affective states occurring during WD can directly lead to relapse after periods of abstinence [41-43]. Our previous studies demonstrated that chronic nicotine WD elicits anxiogenic effects [14,15,26] that are precluded by systemic pharmacological blockade of ERBB4 receptors [15]. In this study, we used genetic, biochemical, and functional approaches to investigate NRG3-ERBB4 neural correlates within the brain anxiety circuitry during chronic nicotine exposure and WD. At the cellular level, both *Nrg3* and *ErbB4* mRNAs increased in the VH at 24hWD. Moreover, attenuation of anxiety-like phenotypes in VH *ErbB4* KD mice was accompanied by alterations in inhibitory transmission and overall network activity in the CA1 area, a subregion of the VH enriched in *Nrg3* and *ErbB4* mRNAs, during 24hWD from chronic nicotine. Thus, our findings determined that NRG3 and its cognate receptor ERBB4 are necessary for the manifestation of anxiety-like behavior during nicotine WD, a hallmark symptom seen in mice [44,45] and humans [4,46,47]. Therefore, in agreement with our previous studies [15,22], our data provide compelling evidence that the NRG3-ERBB4 signaling pathway represents a causal and druggable mechanism in the expression of nicotine WD phenotypes.

### Synaptic expression of ventral hippocampal NRG3 and ERBB4 are induced during 24hWD

Active gene transcription is necessary for addiction processes. However, divergent gene networks may underlie various addiction phenotypes. As mentioned previously, CREB is a well-established transcription factor in the field of addiction [48-50]. It is crucial for stimulus-transcription coupling, in which events occurring at the cell surface leads to alterations in gene expression and ultimately changing neuronal protein expression. Our lab has previously demonstrated that increases in total and phosphorylated levels of CREB are present during chronic nicotine and 24hWD in the hippocampus, leading to increased transcription of the CREB target gene, *Nrg3* [15]. In the present study, we detected increased *Nrg3* transcripts specifically at the 24hWD time point within the VH, with 1 week post WD mRNA levels of *Nrg3* returning to baseline. Interestingly, this increase in *Nrg3* transcription falls within the timeline of observable nicotine WD signs in rodents—being the most severe 24-48 h post-WD and tapering off after this timepoint [51]. Similarly, studies in smokers show that WD symptoms set in between 4-24 hours after an individual smokes their last cigarette, with the symptoms peaking at around day 3 [52]. Moreover, smokers exhibit functional and morphological changes in the hippocampus during abstinence [53,54].

24hWD-induced molecular changes were also seen at the protein level, with increased NRG3 and ERBB4 protein levels in synaptic fractions. Previous studies have demonstrated synaptic enrichment of these proteins, where they participate in the formation and maintenance of synapses [21,33,55,56]. Western blotting of VH synaptosomal fractions showed the presence of the full-length NRG3 (75kDa) [57] and full-length ERBB4 (180kDa) along with cleavage products (120kDa and 80kDa bands) [34,58], all of which were upregulated at 24hWD (Fig.1B). NRG binding is implicated in the shedding of a 120kDa ectodomain fragment via cleavage by the metalloprotease TNF-α-converting enzyme (TACE) [34]. Subsequent cleavage by presenilin/gamma secretases releases an 80kDa intracellular domain with an active tyrosine kinase domain and can translocate to the nucleus and promote nuclear transcription of various transcription factors [59,60]. ERBB4 appears to be the only erbB that harbors a nuclear localization signaling in its intracellular domain [61]. Our data suggest that 24hWD not only induces translation of the full-length precursor ERBB4 further but provokes proteolytic cleavage as well (Fig.1Bii-iv). Further experimentation is necessary to determine the precise role of nuclear ERBB4 signaling during WD.

### Ventral hippocampal ERBB4 signaling mediates WD-induced anxiety-like behaviors

We found that increased expression of NRG3-ERBB4 signaling in the VH corresponds to anxiety-like behaviors, with genetic disruption of this pathway eliminating these phenotypes induced during 24hWD. The NIH test and the Open Field exploratory tests model ascertains anxiety-like behaviors in rodents by integrating an approach-avoidance conflict paradigm [62]. Our behavioral data is in accordance with our previous studies [15,24-26,63], revealing anxiety-like responses increase during 24hWD. These findings suggest that abstinence from nicotine is an additional stressor that promotes anxiogenic behaviors when placed in an unfamiliar environment, a phenotype detected in both the NIH and OF tests.

The OF test revealed that VH *ErbB4* deletion has an anxiolytic effect on 24hWD-dependent exploratory behavior in an unfamiliar environment. Yet, the NIH test demonstrated a significant baseline difference in the latency to feed in VH *ErbB4* KD mice across all treatments on Novel Test Day. These findings suggest that *ErbB4* deletion produces a floor effect in this paradigm, where anxiety-like responses are at such a low threshold that no treatment effects are detectable. Baseline levels of hyponeophagia in mice can be influenced by genetic background, isolated housing, and specific genetic mutations that affect anxiety-related behaviors [64,65]. Approach-avoidant paradigms are common exploration-based tests for anxiety-like behaviors, based on the premise that novel environmental stimuli may be perceived as threatening and thereby inhibit the innate tendency to explore. The NIH and OF assays both rely on response to novel stimuli, therefore our data may alternatively indicate alterations in novelty-seeking phenotypes rather than the described impact on avoidance. Nonetheless, altered novelty-seeking and harm avoidance are both putatively linked to increased risk for drug abuse, as well as anxiety and related disorders [66-69].

Current literature examining pharmacological and genetic manipulations of NRG3 and ERBB4 signaling in the brain result in a similar display of abnormal affective behaviors in mice, strengthening our evidence that *ErbB4* KD impacts anxiety-related behavior. In adult mice, neonatal overexpression of NRG3 results in increased impulsive action, heightened anxiety, and reduced social function [70]. *ErbB4* KD models report that deletion of *ErbB4* from interneurons in the cortex and hippocampus results in decreased anxiety [71-73]. Conversely, it seems this signaling pathway may mediate opposing behaviors in other brain regions. For example, *ErbB4* deletion from serotonergic neurons in the dorsal raphe nucleus engenders anxiogenic behaviors, which is reversed by the inactivation of this subset of neurons [74]. In the amygdala, deletion of *ErbB4* from somatostatin (SOM+)-expressing interneurons also increases anxiety [75], whereas administration of NRG1 alleviates anxiety and enhances GABAergic transmission [76]. Additionally, blocking NRG1-ERBB4 signaling in the bed nucleus of the stria terminalis (BNST) region had anxiogenic effects on behavior as well [77]. These data provide evidence of brain region and circuitry-specific modulation of NRG-ERBB4 signaling on select phenotypes. Recent studies have shown that ventral CA1 cell projections, the primary output of the hippocampal formation, mediate anxiety-like behavior via reciprocal communication with the medial prefrontal cortex (mPFC) [78-82], hypothalamus [83], and amygdala [84,85]. Therefore, these findings support the hypothesis that *ErbB4* deletion could attenuate the effects 24hWD has on VH specific function and its associated circuitry.

### Ventral hippocampal ErbB4 knock-down mice have decreased sIPSC and mIPSC frequencies and dysregulated network clustering in the CA1 area during 24hWD

The impact of VH *ErbB4* deletion on pyramidal cell dynamics during 24hWD is unknown. smFISH experiments demonstrated the highest expression of *ErbB4* mRNA in the CA1 area. In addition to pyramidal cells, there are over 20 types of GABAergic inhibitory interneurons in the CA1 subregion alone [86]. ERBB4 protein is particularly enriched in postsynaptic terminals of parvalbumin (PV+) and cholecystokinin (CCK+)- expressing GABAergic interneurons within the hippocampus [87-89]. Within the hippocampus and the cortex, NRG3 expression is restricted to excitatory presynaptic pyramidal cell terminals contacting interneurons, promoting formation and maturation of excitatory synapses onto ERBB4+ interneurons [33,55,90]. ERBB4 regulates GABAergic neurotransmission, synaptic plasticity, and neuronal activity in cortical areas [88,91,92]. In our study, GABAergic synaptic transmission was weakened in VH *ErbB4* KD mice during 24hWD, characterized by a significant reduction in the frequency of sIPSCs and mIPSCs recorded from pyramidal neurons. A similar IPSC frequency decrease has been previously reported after *ErbB4* deletion in fast-spiking interneurons [71]. Changes in synaptic current frequency, particularly at the level of action-potential independent mIPSCs, are traditionally attributed to pre-synaptic mechanisms, suggesting deficits at the level of GABA releasing interneurons. However, while we see no significant changes in sIPSC amplitude, a measure reflecting post-synaptic GABA_A_ receptor activation, we do find an increase in the amplitude of mIPSCs, as shown after *ErbB4* deletion in the amygdala [93]. It is possible that post-synaptic receptor up-regulation compensates for reduced GABA release after *ErbB4* KD in the VH. Given that mIPSC data isolates receptor responses at individual synaptic boutons, differences in the effect of *ErbB4* KD on sIPSC and mIPSC amplitude may reflect additional adaptations at the level of synaptic arborization following nicotine withdrawal.

Furthermore, the attenuation of mIPSC frequency was accompanied by altered network clustering, suggesting a role for ERBB4 receptors on pyramidal cell ensembles through GABAergic transmission. For example, deletion of the *ErbB4* gene reduces GABAergic transmission, increases the firing of pyramidal neurons, and enhances long-term potentiation (LTP) in brain slices [91,94-96]. Additionally, recent evidence showed that NRG3 strengthens excitatory synaptogenesis onto ERBB4-expressed GABAergic neurons and its functionality in the hippocampus [33]. In the present study, we have not assessed whether ERBB4-induced GABAergic neurotransmission during nicotine WD is NRG3-dependent. Nevertheless, our current and previous findings indicating that nicotine exposure and WD increase gene and protein expression of NRG3, but not NRG1, are consistent with this hypothesis [15].

The electrophysiology and Ca^2+^ imaging experiments were designed to evaluate the effect of VH ErbB4 KD on synaptic transmission and overall network excitability during 24h WD. Previous data from our lab has examined the impacts of both nicotine and 24hWD on VH activity [26]. Using voltage-sensitive dye imaging, CA1 responsivity to schaffer collateral stimulation is increased following *in vivo* chronic nicotine treatment and returns to saline levels after 24h of nicotine cessation; however, GABAergic tone remained disrupted following nicotine withdrawal [26]. Therefore, in this study we focused on cell-selective molecular mechanisms potentially contributing to the disinhibition observed during withdrawal. These data indicate that ERBB4 disruption decreases inhibitory neurotransmission, likely resulting in reduced disinhibition of the CA1. This is in line with the proposed compartmentalization of ERBB4-NRG3 signaling to hippocampal interneurons.

### Conclusion

Our findings demonstrate contributions of VH specific ErbB4 signaling in anxiety-related behaviors seen during nicotine WD. Mechanistically, NRG3-ERBB4 signaling disruption in the VH attenuates nicotine-induced WD anxiogenic behaviors by altering GABAergic modulation of CA1 pyramidal cell activity. These findings suggest that hindering NRG3 activation of ERBB4 at GABAergic synapses curtails inhibitory inputs and regulation of excitatory pyramidal cell activity, and subsequent net output to other brain areas within the anxiety circuitry. Our findings link neuronal mechanisms and circuit-specific effects of NRG3-ERBB4 signaling during nicotine and WD to anxiety-like behaviors, suggesting that targeting NRG3-ERBB4 pathway may advance the development of personalized therapies to smoking cessation.

## Methods and Materials

### Murine subjects

Male and female *ErbB4*-floxed mice (strain B6;129-*ErbB4*tm1Fej/Mmucd, stock number 010439-UCD) were cryo-recovered by the Mutant Mouse Resource and Research Centers (MMRRC), University of California, Davis. Live animals bred in house were 6-8 weeks of age at the beginning of experimentation. Mice were maintained on a 12 h light-dark cycle (lights on at 7:00 AM), with ad libitum food and water. All behavioral procedures were conducted during the hours of 9:00 AM – 5:00 PM. Protocols regarding the proposed work for these studies is approved by the University of Kentucky Institutional Care and Use Committee (IACUC) and the Institutional Biosafety Committee (IBC).

### Stereotaxic surgery and ventral hippocampal microinjections

Surgery was performed on adult mice (6-8 weeks old). After induction of anesthesia with isoflurane (4%), mice were secured in a stereotaxic frame (Stoelting, Wood Dale, IL.). Mice were maintained under isoflurane anesthesia (1-2%) throughout the surgical procedure via a nose cone. Holes were drilled bilaterally into the skull at the injection sites. Ventral intrahippocampal stereotaxic coordinates were measured from the skull surface as follows: AP -2.9, ML ±3.0, DV -3.8. A 33-gauge needle attached to a 5 µl Hamilton syringe was mounted to the stereotaxic frame and, under control of a KDS310 Nano Pump (KD Scientific, Holliston, MA), was used to inject 0.5 µl of 1 × 10^9^ gc/µl AAV at each site. Injections occurred at a rate of 0.1 µl/min, after which the needle was left in place for an additional 4 min. After injections were completed, the skin was sutured, and animals were given an intraperitoneal injection of 5 mg/kg meloxicam (Metacam, Boehringer, St. Joseph, MO) and allowed to recover for up to 1 h on a heating pad before being returned to their home cage. Mice remained in their home cage for an additional 4 weeks until the beginning of novelty-induced hypophagia (NIH) training. Reverse transcriptase coupled quantitative PCR (RTqPCR) analyses of *ErbB4* knock-down (KD) was assessed following all experiments and mice with less than a 20% KD of *ErbB4* in the VH were excluded from experiments.

### Drugs and administration

(-)-Nicotine tartrate (MP Biomedicals, Solon, OH) was dissolved in 0.9% saline. Nicotine was administered subcutaneously via osmotic minipumps (Alzet model 2002, Cupertino, CA) at a dose of 12 mg/kg/d for 14 days, calculated based on the daily pump rate of the pulsatile delivery system (see “pulsatile delivery” below). This dose, reported as freebase weight and based on previous work [15,24-26], corresponds to plasma levels of ∼0.2 μm [27], a concentration within the range observed in human smokers consuming an average of 12 cigarettes a day (plasma levels between 0.04 and 0.21 μm) [27].

### Osmotic minipumps surgeries

#### Pulsatile delivery

A pulsatile nicotine delivery system was achieved by attaching osmotic minipumps to polyethylene (PE60) tubing, similar to that described in Brynildsen and colleagues [28]. The PE60 tubing was prepared using a coiling technique, which consisted of coiling the tubing around a cylinder with a similar circumference to the minipump and dipping the thermoformable tubing in hot water, followed by immersion in ice-cold water. This shaped the tubing into a coil for easy subcutaneous implantation. Once formed, the PE60 tubing was filled with alternating 0.5 µl volumes of nicotine tartrate (or saline for controls) and mineral oil. The model 2002 osmotic minipumps used for experimentation have a delivery rate of 0.5 µl/hr, therefore we developed a 1 hr “on”, 1 hr “off” pulsatile nicotine delivery system. The attached PE60 tubing was filled to a volume that ensured a time course of 14-day intermittent treatment.

#### Minipump treatment groups

In all experiments, animals were implanted with osmotic minipumps to deliver pulsatile administration of either nicotine (12 mg/kg/day) or saline. Following 2 weeks of chronic administration, mice were anesthetized with an isoflurane/oxygen vapor mixture (1–3%), an incision was made above the pump at shoulder level, and the pump was either removed (to initiate a spontaneous WD from either nicotine or saline) or left in place (to serve as sham surgical controls in the nicotine and saline groups). The incision was then closed with 7 mm stainless steel wound clips. Animals were allocated to the different experimental groups based on sex, pre-operative weight average, and behavioral baseline measures.

### Adeno-associated virus production

The University of Pennsylvania Vector Core generated neuron-selective AAV serotype 9 for expressing: Cre recombinase (AAV-CRE: AAV9.CMV.PI.Cre.rBG, titer 1.644 × 10^13^ genome copies (gc)/ml), red fluorescent protein (AAV-RFP: AAV9.CMV.TurboRFP.WPRE.rBG, titer 32.87 × 10^13^ gc/ml), and GCaMP6f Ca^2+^ indicator (AAV-9-PV2822: AAV9.Syn.GCaMP6f.WPRE.SV40). Purification of the vectors was performed using CsCl sedimentation, and vector gc quantification was performed by the UPenn Vector core using qPCR. AAVs were diluted in sterile PBS for microinjections directly into the VH. The University of Kentucky animal facility is equipped to handle appropriate care of animals infected with AAV and all procedures are approved by IACUC. All procedures utilizing AAV are also approved by our Institutional Biosafety Committee.

### Novelty-induced hypophagia (NIH) test

The NIH test was performed as previously described [25]. Briefly, NIH training and testing consisted of exposing mice to a highly palatable food (Reese’s peanut butter chips (Nestle, Glendale, CA (ingredients: partially defatted peanuts, sugar partially hydrogenated vegetable oil, corn syrup solids, dextrose, reduced minerals whey, salt vanillin, artificial flavor, soy lecithin)) and latency to consume was measured. One week before NIH training and for the duration of the experiment, mice were housed in groups of two. Training consisted of daily sessions in which mice were exposed to Reese’s peanut butter chips in a clear plastic dish. Plastic dividers (dividing the standard mouse cage lengthwise) were placed inside each cage to separate the mice during the training and home cage testing periods. Mice were acclimated to the barriers for 1 h before placement of food. Food was placed in the cage for 15 min, and latency to consume was measured. By the 10th day, a baseline latency to approach and consume the food was reached such that there was <20% variability between mice. After the last training session, the amount consumed was recorded as grams of peanut butter chips to ensure there were no appetitive treatment effects. Following training, mice were implanted with 14-day osmotic minipumps filled with pulsatile nicotine (12 mg/kg/day) or 0.9% pulsatile saline. Testing in the home cage (Home Test Day) and novel environment (Novel Test Day) occurred on the last 2 days of minipump viability. On Home Test Day, following testing, minipumps were surgically removed for the WD groups, and sham surgeries were performed on the chronic nicotine group as well as saline animals. Twenty-four hours later, on the Novel Test Day, mice were removed from the home cage and placed in an empty standard cage with no bedding that had been wiped with a cleanser (Pine-Sol, 1:10 dilution) to emit a novel odor and placed in a white box with bright light illumination (2150 lux). Latency to consume the palatable food was recorded on both days.

### Open Field exploratory test (OF)

The Open Field exploratory test is an anxiety-related behavioral model, which also allows simultaneous assay of overall locomotor activity levels in mice. All mice were tested 24 h after nicotine minipumps were removed from the 24h WD groups and sham surgeries were performed for the nicotine and saline groups. Test chambers were wiped with 70% ethanol in between tests to remove any scent cues left by the previous mouse. The ethanol was allowed to dry completely before each testing, and every testing session lasted for 10 min. For the analysis, Top Scan (Clever Sys Inc., Reston, Virginia) software was utilized to track and evaluate mouse movement. Prior to tracking analyses for each mouse, a background profile was generated, and the testing chamber was calibrated in arena design mode according to the manufacturer’s instructions. Software output for each individual test includes total distance moved (in mm) and the time spent in the center (in %). These data were then normalized to the AAV-RFP saline control group. On both behavioral paradigms, the data were analyzed by an investigator blinded to the experimental groups.

### Reverse transcriptase coupled quantitative polymerase chain reaction (RTqPCR)

RTqPCR was performed as previously described [29] on VH or DH samples across all treatment groups. Briefly, RNA was isolated using the RNeasy Mini kit (Qiagen, Hilden, Germany), and 500 ng of RNA was reverse transcribed into cDNA using Oligo dT primer (Promega, Madison, WI) and Superscript II reverse transcriptase (Invitrogen, Waltham, MA). qPCR reactions were assembled using Thermo Scientific Maxima SYBR Green master mix along with 100 nM primers (Eurofins Scientific, Luxembourg). The mRNA levels were determined using the 2^-ΔΔCT^ method [30], and target genes were normalized to the housekeeping gene Hypoxanthine Phosphoribosyltransferase (HPRT). All gene expression values were normalized to their respective AAV-RFP saline-treated controls. Primer sequences are shown in Table S1.

### Synaptosomal preparation

To obtain synaptosomes, frozen VH tissue was weighed and gently homogenized with a glass dounce tissue grinder in 10 vol (1:10, wt/vol) of Syn-PER synaptic protein extraction reagent (Thermo Fisher Scientific, Rockford, IL) supplemented with a protease and phosphatase inhibitor cocktail. Following manufacturer’s instructions, the homogenate was centrifuged at 1200 xg for 10 min at 4°C, and then the supernatant was centrifuged for a further 20 min at 1500 xg at 4°C. The supernatant (cytosolic fraction) was removed, and the synaptosome pellets were resuspended in Syn-PER reagent. The protein concentrations of synaptosomal and cytosolic fractions were determined by the BCA method (Thermo Fisher Scientific).

### Western blotting

Protein analysis was performed as described previously on VH samples of all treatment groups. Briefly, 20 μg of protein was resolved in AnykD™ precast polyacrylamide gel (Bio-Rad Laboratories Inc., Hercules, CA) and transferred to nitrocellulose membranes. Membranes were incubated with LI-COR blocking buffer (LI-COR, Lincoln, NE) for 1 h at room temperature before reacting overnight at 4°C with primary antibodies: Neuregulin-3 (NRG3) (1:500, sc-67002, N-terminal extracellular domain, Santa Cruz Biotechnology, Santa Cruz, CA), ERBB4 (1:500, sc-283, Santa Cruz Biotechnology), and Beta-tubulin (1:2000, 2128L, Cell Signaling Technology, Danvers, MA). After washing in phosphate-buffered saline-Tween-20 (PBST), the blots were incubated in fluorescent secondary antibodies IRDye 800CW Goat anti-Mouse (1:20000, LI-COR) and IRDye 680LT Goat anti-Rabbit (1:20000, LI-COR) diluted in LI-COR blocking buffer for 1 h at room temperature. Membranes were then washed, and immunolabeling detection and densitometry measurements were performed using the LICOR Odyssey System (LI-COR). Ratios of the proteins of interest (NRG3 and ERBB4) to the housekeeping protein (β-tubulin) densities were calculated for each sample and normalized to AAV-RFP saline-treated controls. The same Beta-tubulin control bands were used to calculate ErbB4 protein content on blots containing all three densities of ErbB4 resulting from a single probe.

### Stellaris single-molecule fluorescent in situ hybridizations (smFISH)

Forty-eight antisense ‘Stellaris probes’ oligonucleotide probes for mouse *Nrg3* and *ErbB4* were designed using Biosearch custom design algorithms and synthesized with a 5’ Quasar 570 and 670 labels, respectively. One brain hemisphere from each mouse was collected and fixed overnight (4°C) in sterile 2% paraformaldehyde solution prepared in PBS. Fixed brains were cryoprotected in 15% sucrose overnight, followed by 30% sucrose overnight incubation (4°C). Cryoprotected brain hemispheres were horizontally sectioned through the VH at 30 µm and processed for FISH experiments. FISH was performed as described previously [31] with few modifications as follows. Slides were brought to room temperature, and all steps were performed at room temperature unless indicated otherwise. Warmed tissue sections were washed three times with 20 mM Glycine in 1X PBS 5 min each, followed by 3 washes in freshly prepared 25 mM NaBH_4_ in 1X PBS, 5 min each. After a quick rinse with 0.1 M TEA, slices were washed in a 0.1 M TEA + 0.25% acetic anhydride solution for 10 min, followed by a 3 min wash in 2x SSC. Slices were then dehydrated in 70, 95, and 100% EtOH (3 min each), and immediately de-lipidized in chloroform for 5 min and rehydrated. Next, sections were washed 2 times in 2x SSC, followed by a quick wash in 0.3% triton X-100, before hybridization buffer was applied. *Nrg3* probes, *ErbB4* probes, and scrambled control probes were resuspended in TE buffer to a final concentration of 25 µM and added to hybridization buffer at 1:100 dilution. Hybridization was performed for 12-16 h at 37°C. Samples were then coverslipped the following day using Prolong Gold mounting medium with DAPI stain (Invitrogen) and analyzed by epifluorescent microscopy. Leica DMI6000 epifluorescent microscope with ORCA Flash ER CCD camera (Hamamatsu, Japan) was used for imaging unless otherwise specified. For quantification between samples, imaging parameters were matched for exposure, gain, offset, and post-processing. Scrambled probes were used as a control, and to assign image acquisition parameters that would minimize any nonspecific signal from the scrambled probe. *Nrg3* and *ErbB4* mRNA particle numbers were quantified using ImageJ and normalized to the number of nuclei. For each subregion (DG, CA3, CA1) of the VH, three images were taken, quantified, and averaged to represent an n of 1.

### Whole-cell patch-clamp electrophysiology

24 h after minipump removal and immediately after behavioral testing, mice were sacrificed via live decapitation, brains were rapidly removed, and horizontal slices (300 μm-thick) containing the ventral hippocampus were cut using a vibratome (VT1200S; Leica Microsystems, Wetzlar, Germany) in an ice-cold cutting solution, containing the following (in mM): 93 NMDG, 2.5 KCl, 1.25 NaH_2_PO_4_, 30 NaHCO_3_, 20 HEPES, 25 glucose, 5 Na-ascorbate, 2 thiourea, 3 Na-pyruvate, 10 MgSO_4_, and 0.5 CaCl_2_, adjusted to 300–310 mOsm, pH 7.4 and continuously oxygenated with 95% O_2_ and 5% CO_2_. Slices were allowed to recover in the cutting solution at 34–36°C for 30 min and were thereafter maintained in an oxygenated recording artificial cerebrospinal fluid (aCSF) solution at room temperature. Recording aCSF contained the following (in mM): 130 NaCl, 3 KCl, 1.25 NaH_2_PO_4_, 26 NaHCO_3_, 10 glucose, 1 MgCl_2_, and 2 CaCl_2_, pH 7.2–7.4, when saturated with 95% O_2_ and 5% CO_2_.

For electrophysiology recordings, recording pipettes were pulled from borosilicate glass capillaries (World Precision Instruments, Sarasota, FL) to a resistance of 4–7 MΩ when filled with the intracellular solution. The intracellular solution contained the following (in mM): 145 potassium gluconate, 2 MgCl_2_, 2.5 KCl, 2.5 NaCl, 0.1 BAPTA, 10 HEPES, 2 Mg-ATP, and 0.5 GTP-Tris, pH 7.2–7.3, with KOH, osmolarity 280–290 mOsm. Brain slices were transferred to the recording chamber continually perfused with carbogen-saturated recording aCSF (1.5-2.0 ml/min) at 31–33°C. Pyramidal neurons in the CA1 region were viewed under an upright microscope (Olympus BX51WI) with infrared differential interference contrast optics and a 40x water-immersion objective. To evaluate spontaneous inhibitory postsynaptic currents (sIPSCs), the cells were voltage-clamped at 0 mV; to evaluate spontaneous excitatory postsynaptic currents (sEPSCs), the cells were voltage-clamped at -70 mV. To isolate miniature excitatory postsynaptic currents (mEPSCs) and miniature inhibitory postsynaptic currents (mIPSCs), recording aCSF included 1 µM tetrodotoxin.

Currents were low pass filtered at 2 kHz and digitized at 20 kHz using a Digidata 1550B acquisition system (Molecular Devices, San Jose, CA) coupled to the Multiclamp 700B amplifier (Molecular Devices) and pClamp11 software (Molecular Devices). Access resistance (10–30 MΩ) was monitored during recordings by injection of 10 mV hyperpolarizing pulses; data were discarded if access resistance changed >25% over the course of data collection. All analyses were completed using Clampfit 11.1 (Molecular Devices).

### Calcium imaging

Two-minute videos of Ca^2+^ fluorescence were acquired with an ORCA-Flash 4.0 (V2) digital camera (Hamamatsu) via a 40x objective at 25 frames/second with a 512×512 pixel binning. Following imaging of spontaneous Ca^2+^ transients, fluorescent signals in response to extracellular stimulation were recorded in each field of view. Extracellular stimuli were delivered by 100 μs current pulses generated by a Master-9 stimulator (A.M.P.I) via a bipolar tungsten electrode positioned just outside the field of view in the stratum radiatum. The amplitude of the current pulses was controlled by a stimulus isolation unit (ISO-Flex, A.M.P.I.). Minimal stimulation was defined as minimal current intensity to produce visible fluorescence in any of the imaged cells. Network analysis of Ca^2+^ transients, was performed on the basis of undirected adjacency matrices thresholded to preserve 25% of the strongest correlation coefficients as described in our previous publication [32].

### Data analyses

Statistical analyses were performed with GraphPad Prism 6.0 software package (GraphPad Software, San Diego, CA). Protein and mRNA analysis was performed using ordinary one-way or two-way ANOVA. Behavioral measures, except where noted, were analyzed using two-way repeated-measures (RM) ANOVA as test day or genotype as within-subject variables and drug treatment as a between-subject variable. Network analyses of Ca^2+^ transients were done with one-way or two-way RM ANOVA, as indicated. ANOVA was followed by Sidak’s multiple comparison tests. For the electrophysiological studies, amplitudes of EPSCs and IPSCs were computed from an average of 50–100 individual current traces. Mean EPSC and IPSC frequencies were analyzed from 20 s long trace segments. EPSC and IPSC data were analyzed using 2-tailed Student’s t-tests. All data were expressed as mean ± SEM, and significance was set at p<0.05.

## Author Contributions

M.L.F. and J.R.T. contributed to the conception of the presented idea. M.L.F., T.A., P.S., J.L.T., P.I.O., and J.R.T. planned the experiments. M.L.F., E.R.P., B.O., T.A., P.S., J.L.T., and P.I.O. ‘carried out the experiments. All authors contributed to data analysis. M.L.F., E.R.P., P.I.O., and J.R.T. wrote the manuscript.

## Supplemental Materials

**Figure S1.**
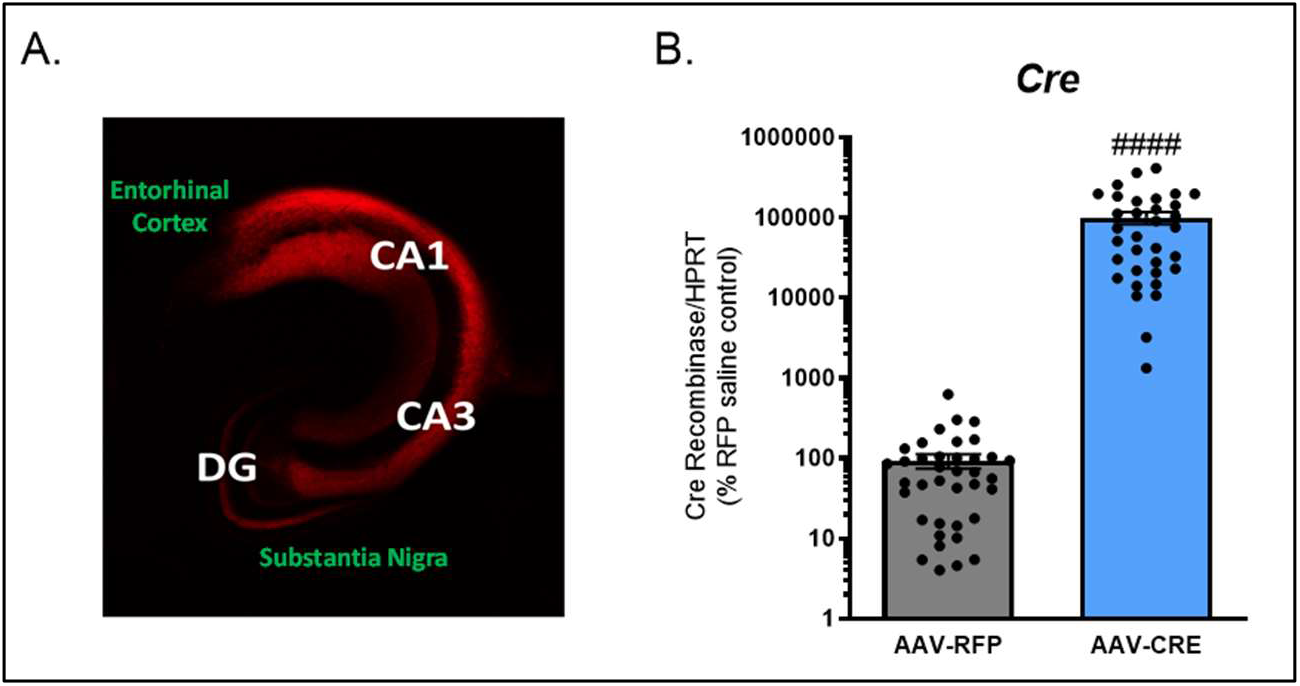
Viral expression analysis of VH CRE recombinase. (A) Representative con-focal image of RFP in VH (white) and non-expressing (green) regions. (B) Quantification via RTqPCR analyses of CRE recombinase mRNA in RFP- and CRE-injected mice (####p<0.0001).

**Table S1.**
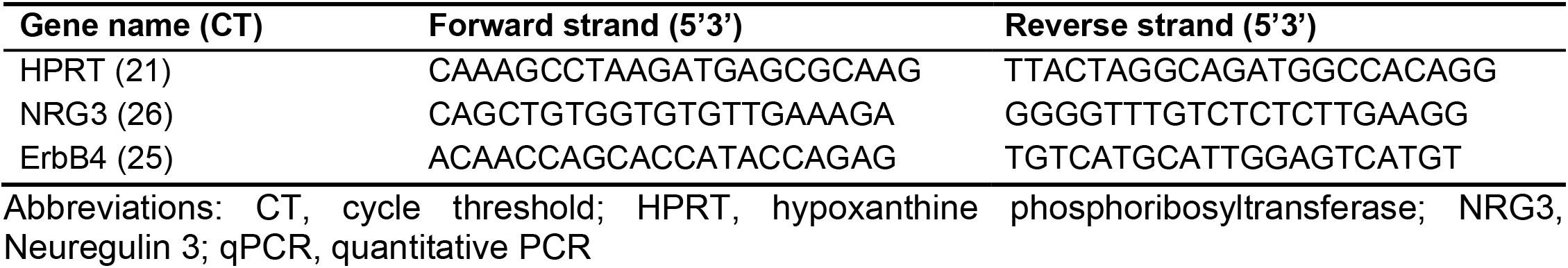
RTqPCR Primers

